# A Modified Approach to Define Walking Center of Mass Mechanical Energy Recovery: Human Walking Involves Energy Loss Throughout Stance

**DOI:** 10.1101/2024.07.09.602732

**Authors:** Seyed-Saleh Hosseini-Yazdi, John EA Bertram

**Author notes:** **Correspondence Address:** John EA Bertram, Biomedical Engineering Department, Schulich School of Engineering, University of Calgary, 2500 University Dr NW, Calgary AB T2N1N4. **Contributions:** Both authors contributed to concept development, analytical modeling and empirical analysis, writing, and editing.

## Abstract

An exchange between potential and kinetic energy over the step has long been considered a key feature in the energetic effectiveness of human walking. However, it is difficult to identify mechanisms responsible for limiting such an exchange in human walking. This study proposes a modified definition of center-of-mass (COM) energy recovery (*R*_*c*_) that quantifies the proportion of mechanical energy transferred from one step to the next while accounting for total step dissipation. Simulations show that *R*_*c*_ decreases nearly linearly with walking speed on level ground, indicating no preferred speed. This behavior arises from analytical formulations that neglect active work during single support (pendular motion). In contrast, empirical data reveal consistently lower *R*_*c*_, likely due to elevated collision losses or negative net single-support work not captured by the analytical model. When both single- and double-support phases are considered analytically, *R*_*c*_ exhibits a maximum of 59.4% at 1.21 m.s^−1^, coinciding with minimal active muscle work over the step. We further show that the *R*_*c*_ trajectory is asymmetric, contrary to prior assumptions, and is governed by total step dissipation. Accordingly, challenging walking conditions associated with higher metabolic cost, such as restricted visual lookahead, are predicted to reduce *R*_*c*_ (maximum 58.5%).

## Introduction

There would be no mechanical energy required if human walking operated as a true inverted pendulum with sufficient initial momentum (Alexander, 1992b, 1995). However, human walking involves some mechanical energy loss (Donelan et al., 2002a) and only a portion of the mechanical energy remains available for the next step. The Center of Mass (COM) mechanical energy recovery has been defined based on potential (*E*_*p*_) and kinetic (*E*_*k*_) energy exchange during COM pendular motion (Cavagna et al., 1976). Theoretically, if the exchange is perfect, no net mechanical work is required. However, when the exchange is incomplete, some mechanical energy must be supplied by muscular work Cavagna et al.,1977). Therefore, Cavagna (Cavagna et al., 1977) defined the pendular motion mechanical energy recovery (*R*_*IP*_) as the fraction of mechanical energy that appears conserved through the exchange between *E*_*p*_ and *E*_*k*_, rather than being actively generated by muscles. It is a key metric in walking biomechanics that measures potential efficiency of energy transfer between steps, reflecting how well mechanical energy can be conserved during gait. A higher *R*_*IP*_ indicates less active mechanical work and, such, reduced metabolic cost (Cavagna et al., 1977; Donelan et al., 2002a). Since the original definition of *R*_*IP*_ is based on the premise of pendular motion (Cavagna et al., 1976, Cavagna et al., 1977), it does as consider the separate influences of mechanical work for the individual limbs during the step to step transition (Donelan et al., 2002b). We assume that positive external limb work offsets energy dissipation (Kuo, 2002) and maintains the system’s total mechanical energy. Hence, considering total step dissipation (Hosseini-Yazdi & Bertram, 2025b; Ruina et al., 2005) might present a new aspect of human gait under varying conditions, such as when the view horizon (lookahead) changes (Hosseini-Yazdi Seyed-Saleh, 2024). Thus, we believe a collisional or energy dissipation approach (Bertram & Hasaneini, 2013; Hosseini-Yazdi & Bertram, 2025b; Ruina et al., 2005) may present a new perspective on COM mechanical energy recovery.

Human locomotion is hypothesized to mimic pendulum and spring-like mechanics to minimize required active muscle work (Alexander, 1995; Kuo, 2002). Specifically, the single support phase of walking is suggested to align with the motion of an inverted pendulum (Kuo et al., 2005). In an idealized scenario with no energy dissipation, a walker starting with sufficient initial energy would require no external mechanical work input. Thus from this perspective, human walking primarily involves converting potential and kinetic energy (Alexander, 1992b, 1995).

According to Cavagna et al. (Cavagna et al., 1976, Cavagna et al., 1977), walkers perform work against gravity (*w*_*v*_) to lift the COM and accelerate it forward (*w*_*f*_). When potential and kinetic energies are completely out of phase, all mechanical energy transitions between gravitational potential energy and kinetic energy. However, when these energies become more aligned, energy exchange is disrupted, requiring active muscle work to maintain consistent speed (*w*_*ext*_). This assumption hinges on the notion that human walking always resembles inverted pendulum mechanics (Figure 1A).

**Figure 1.**
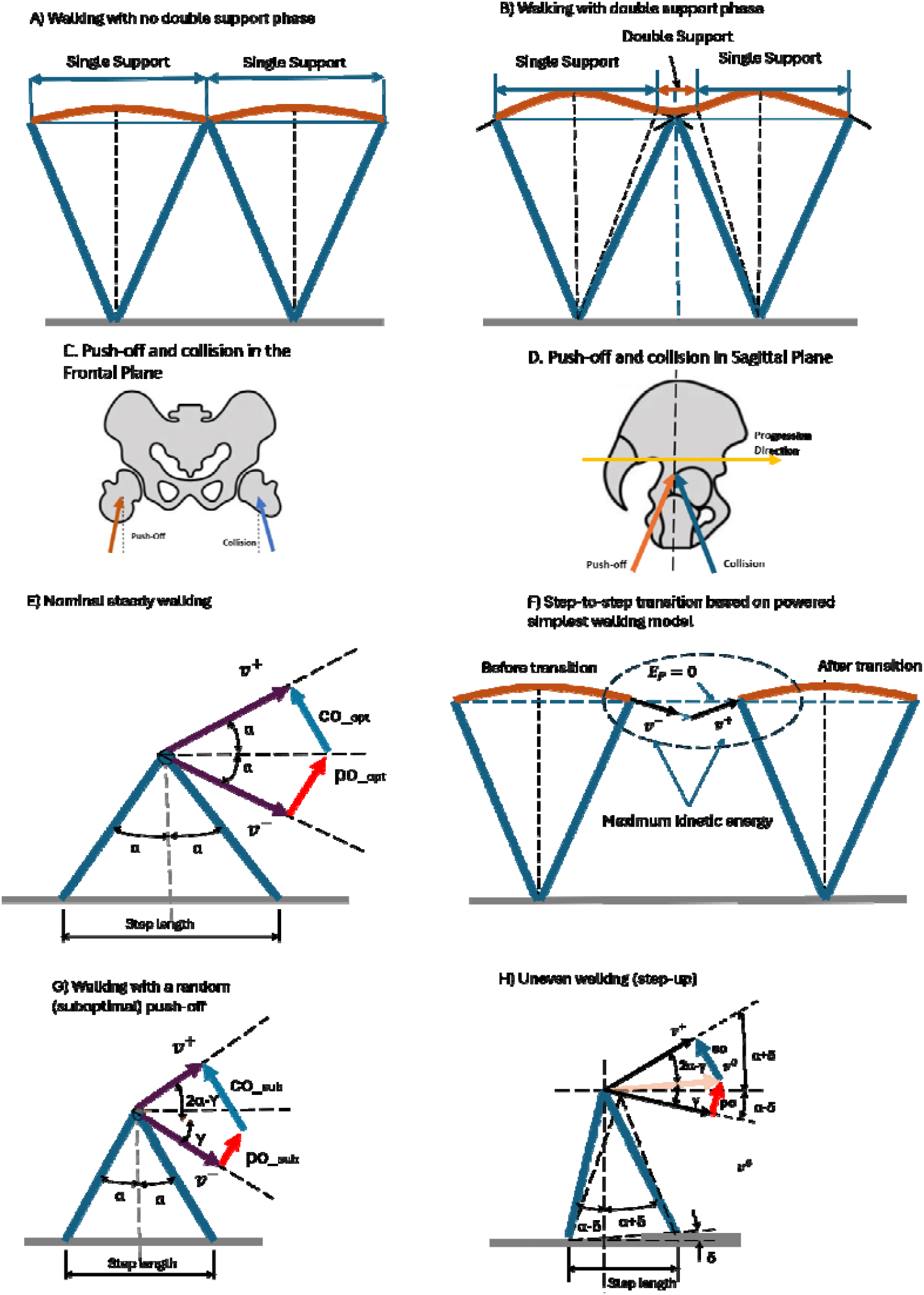
Schematics showing the step-to-step transition during even walking: (A) pendular walking model with rigid legs in which the step-to-step transition is instantaneous (no double support phase), (B) human walking demonstrates a double support phase with a protracted switch from one inverted pendulum to another, (C) push-off and collision impulses have similar effects in the vertical direction, (D) push-off impulse adds to the COM mechanical energy in the progression direction while the collision impulse reduces momentum, (E and F) In nominal steady walking, the push-off and collision work, and pre () and post transition () speeds are equal, the pre transition velocity may be approximated by, (G) if the pre-emptive push-off is less than optimal (suboptimal), the magnitude of the subsequent collision impulse grows and the post transition speed declines, (H) when the total step transition and gravity work are provided by the pre-emptive push-off impulse, optimal uneven walking occurs.

Donelan et al. (Donelan et al., 2002a), suggested that during the step-to-step transition, there is a substantial deviation from pendular motion when COM mechanics transition to the next stance (Figure 1B). They proposed that humans simultaneously perform positive and negative work during step transitions (Donelan et al., 2002b). The positive work is the result of muscle activation to offset step transition energy loss (Kuo, 2002). Post step transition walking tends to resemble pendular motion again (Donelan et al., 2002a; Kuo et al., 2005).

When the legs apply force to the ground surface, they cause changes in the COM path and velocity. A change in the COM velocity is defined as a collision in dynamics. At the transition between stance legs, double support, two events occur that make the transition energetically effective. The previous stance leg exerts a generative collision (push-off) and immediately after the new stance leg produces a dissipative collision (heel-strike) (Ruina et al., 2005). Considering the legs as separate actuators, exerting toe-off (push-off) and heel-strike impulses (Kuo et al., 2005), it becomes reasonable to investigate their separate effects on the COM. Similar to other actuators, the work of each leg is determined by the force applied and the corresponding displacement. Thus, while the net effect on the COM may reflect the resultant force applied, the independent work of each leg incurs its own metabolic cost (Griffin et al., 2003).

Both push-off and heel-strike impulses generate upward motions on the COM in the frontal plane, but their sagittal plane effects differ. The push-off impulse propels the pelvis forward, while the heel-strike impulse opposes forward progression, partially costing momentum and energy (Figure 1C & D) (Donelan et al., 2002a, 2002b; Kuo, 2002). Consequently, the push-off impulse requires muscular work and contributes to forward momentum, whereas the heel-strike impulse results in energy loss as the path of the COM is altered (Donelan et al., 2002a; Kuo, 2002).

A simple powered walking model demonstrates the independent contributions of the leading and trailing legs during step transitions (Kuo, 2002). For a given walking speed, it predicts the active work required (push-off work) for optimal steady-state walking, where active work matches dissipation. A pre-emptive push-off, coming just prior to heel strike, limits the magnitude of the subsequent heel-strike collision (Kuo, 2002; Kuo et al., 2005). However, if the pre-emptive push-off is sub-optimal (Hosseini-Yazdi & Bertram, 2025c; Huang et al., 2015), it can only partially mitigate the resulting collision (Hosseini-Yazdi & Bertram, 2025a, 2025b, 2025c). Hence, to maintain steady walking, additional positive mechanical energy is required during the subsequent single stance phase (Darici & Kuo, 2023; Hosseini-Yazdi & Bertram, 2025a; Kuo, 2002). Therefore, suboptimal push-off (*po*_*sub*_ < *po*_*opt*_) results in increased dissipation and reduced energy transferred to the subsequent step.

Mechanical work required in walking increases with terrain complexity (Hosseini-Yazdi & Kuo, 2025; Voloshina et al., 2013) and age (Das Gupta et al., 2019; Hosseini-Yazdi Seyed-Saleh, 2024). Restricted lookahead might also impair the ability to select appropriate footholds, which could impair walking dynamics (Matthis & Fajen, 2013). It is shown that walking with a restricted lookahead also coincides with increased mechanical work (Hosseini-Yazdi Seyed-Saleh, 2024). Therefore, we can expect lower COM energy recovery.

This study redefines the COM energy recovery (*R*_*c*_) based on following the step energy collision dissipation (Bertram & Hasaneini, 2013). If the entire gait was like an inverted pendulum, the COM energy recovery would be 100%. Since it is also demonstrated that mechanical energy loss may occur in different phases of stance (Hosseini-Yazdi & Bertram, 2025b; Kuo et al., 2005), the potential and kinetic energy exchange may not be the only factor influencing mechanics of the gait cycle (Alexander, 1992a; Geyer et al., 2006). Consequently, active muscle work is to regulate COM mechanical energy. Accordingly, we develop analytical simulations to derive analytical COM energy recovery for nominal (optimal) walking across a range of speeds based on the notion of pendular motion and step transition work. The simulations are extended to scenarios with suboptimal push-off impulses (*po*_*sub*_ < *po*_*opt*_) and cases where COM elevation changes; i.e. step-ups and step-downs or uneven walking. Similar to even walking (Kuo, 2002), we examine a scenario where the pre-emptive push-off provides the entire step transition energy and compensates for work against gravity during uneven walking. Further, we compare the analytical results with *R*_*c*_ derived from empirical COM work trajectories (Hosseini-Yazdi Seyed-Saleh, 2024) for walking over smooth surfaces, tracking the mechanical energy dissipation of different stance phases.

### New Definition for COM Energy Recovery

According to a powered simple walking model (Kuo, 2002), the active push-off (*po*) is applied pre-emptively to the COM to mechanically energize the step-to-step transition (Donelan et al., 2002a). This compensates for the dissipation caused by the heel-strike collision, which occurs after push-off (Donelan et al., 2002b). Consequently, the required external work (*w*_*ext*_) is assumed to offset the energy lost during heel-strike (Cavagna et al., 1977). Considering ‘*u*’ as the transition impulse, its associated work is calculated as 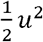 (Kuo, 2002). We count the total mechanical energy of the body as the sum of its stance phase potential and kinetic energies (*E*_*tot*_ = *E*_*P*_ + *E*_*k*_). Assuming the lowest elevation of the COM during stance phase as the reference for gravitational potential energy (datum, *E*_*P*_ = 0), total energy at the end of stance phase corresponds to the maximum kinetic energy (*E*_*K_max*_ = *E*_*tot*_, Figure 1E & F). Contrary to the original inverted pendulum recovery definition (*R*_*IP*_) (Cavagna et al., 1977), we redefine the COM of stance energy recovery based on step work to maintain COM total energy:

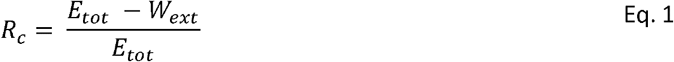

This equation quantifies the proportion of mechanical energy transferred from one step to the next. Since the system’s total energy equals the maximum kinetic energy just before the step transition, the COM energy recovery may be re-written as:

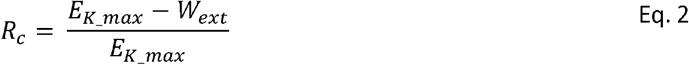

Assuming the COM velocity just before the transition approximates the average walking velocity (*v*_*ave*_ in which 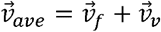) (Kuo, 2002), and *w*_*ext*_ represents the normalized external work per unit mass, the COM energy recovery can be derived as:

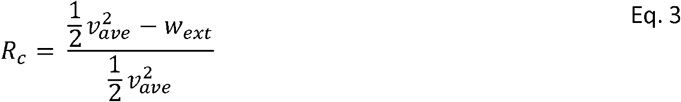

To sustain a steady walk, energy must remain constant before and after the step-to-step transition. Denoting the velocity before and after the transition as *v*^−^ and *v*^+^, respectively, steady walking implies, 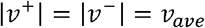. With the kinetic energy conserved across the transition, the active work energy addition (*w*_*ext*_) equals the heel-strike dissipation or collision work (*w*_*co*_). Therefore, the COM energy recovery can be further expressed as:

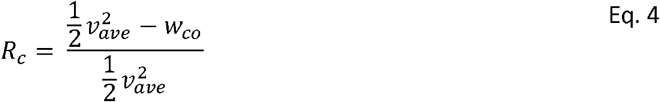

Here, *w*_*co*_ represents the normalized energy loss due to the heel-strike collision.

### Simulation based on a powered simple walking model

In this model, we assume concentrated mass at the pelvis or COM with rigid massless legs. The nominal push-off impulse for a given walking speed is expressed as *po* = *v*_*ave*_ · tan *α* where ‘2*α*’ represents the angle between the leading and trailing legs at the step transition point (Kuo, 2002). Consequently, the nominal push-off work becomes 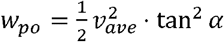 (Kuo, 2002). Assuming equality of step transition positive and negative work (Donelan et al., 2002b), i.e. *w*_*co*_ = *w*_*po*_ or *w*_*ext*_ = *w*_*co*_, the COM energy recovery is:

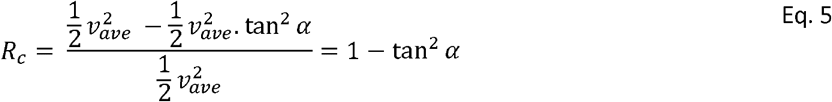

For small angles, the approximation tan *α* ~ *α* holds, leading to *po* = *v*_*ave*_ · *α* and 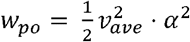. Assuming a unit leg length, the step length (*s*) becomes proportional to 2*α* (Kuo, 2001, 2002). Since step length follows a power law with walking speed (Kuo, 2001); *s* ~ *v*^*0.42*^, yielding 2*α* = *c* · *v*^*0.42*^ and 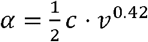. Substituting ‘*α*’ into Eq. 5 with tan *α* ~ *α*:

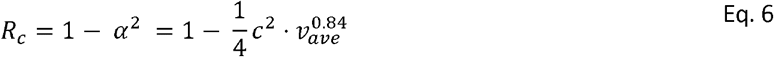

This analytical expression suggests that the COM energy recovery decreases nearly linearly with walking speed. Applying the powered simple walking model to a suboptimal push-off (*po*_*sub*_, Figure 1G) (Hosseini-Yazdi & Bertram, 2025a) the mid-transition velocity (*v*^0^) forms an angle (*γ*) with the COM velocity before transition (*v*^−^) which depends on the induced push-off impulse magnitude: 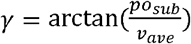. Since *v*^−^ ~ *v*_*ave*_ (Kuo, 2002), the transition collision impulse (*co*) becomes:

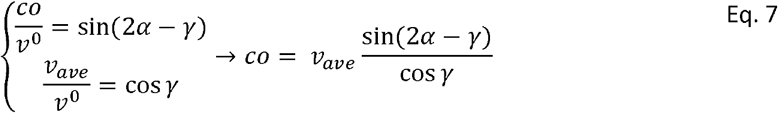

The normalized collision work is *w*_*co*_ = 0.5*co*^2^ (Kuo, 2002), and the modified COM energy recovery is:

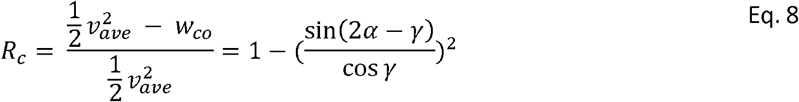

Using small angle approximations for ‘*α*’ and ‘*γ*’:

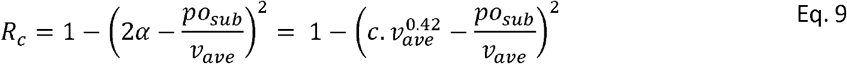

This result shows that the COM energy recovery becomes a nonlinear function of both suboptimal push-off magnitude and average walking speed.

For uneven terrain with a step-up, maintaining average walking speed (*v*_*ave*_) at mid-stance requires the post-transition COM kinetic energy to also account for the work against gravity. The post-transition velocity (*v*^+^) must therefore satisfy (Figure 1H):

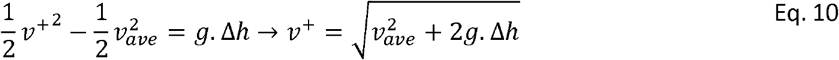

If the entire uneven step energy is provided by a pre-emptive push-off, then:

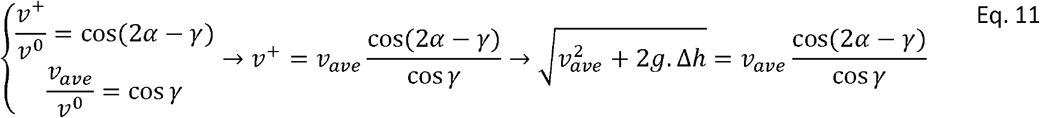

The mid-transition velocity direction angle change (*γ*) must be determined numerically for a given walking condition. The uneven step collision impulse becomes:

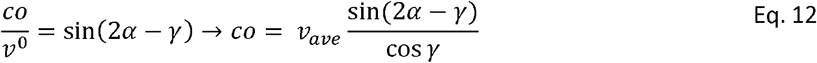

Thus, the COM energy recovery for an uneven step can be derived similar to even walking when a suboptimal push-off is applied, yet with *γ* > *α*. Using the small angle approximation, the COM energy recovery for uneven walking becomes: 1 − (2 *α* − *γ*)^2^. For a step-down, the process is similar, with Δ_*h*_ being negative. Here, the potential energy converts to kinetic energy, increasing momentum, and the required push-off magnitude decreases as the step-down amplitude increases.

### COM energy recovery based on empirical data

For experimental data analysis, the collision work at heel-strike can be estimated by integrating COM power trajectories (Figure 2) (Kuo et al., 2005). Using this approach, the COM energy recovery can be expressed as:

**Figure 2.**
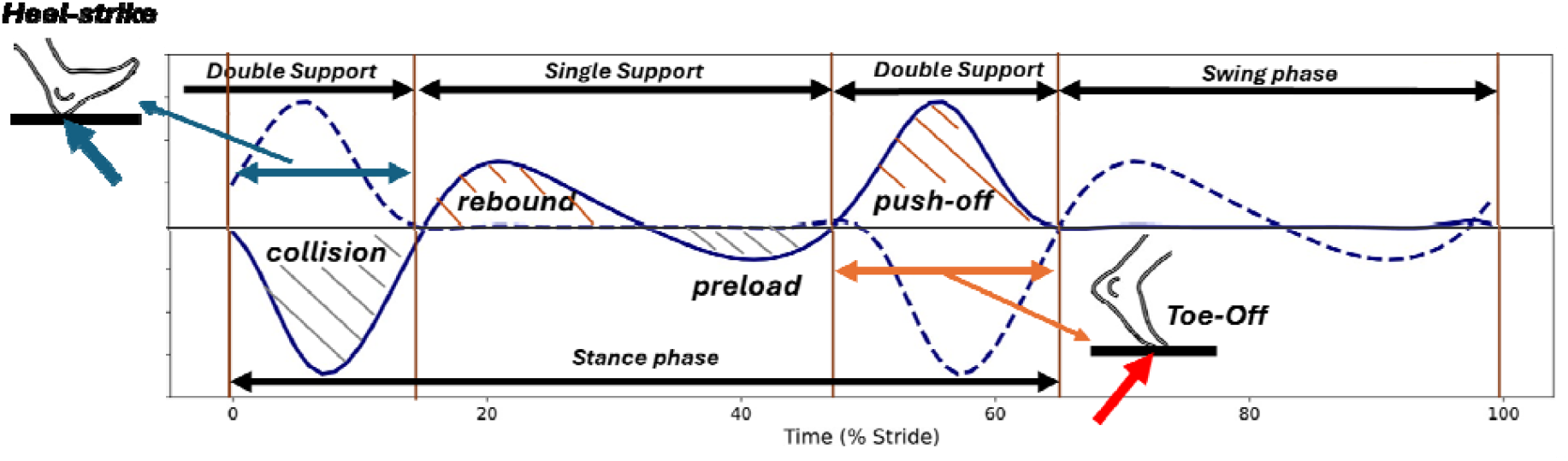
The COM power trajectory during a stride is illustrated, with the power contribution of the opposite leg represented by a dashed line. Each stance begins with a heel-strike, which exerts a collision impulse on the COM. Following the initial double support phase, gait transitions into a single support phase (rebound and preload). In the late stance phase, the trailing leg exerts a push-off impulse during toe-off. The mechanical work for each phase can be determined by integrating the corresponding power trajectory over time.

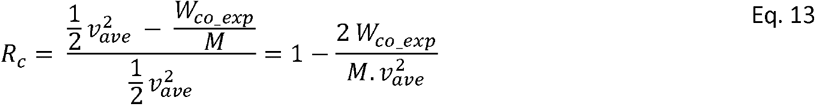

Here, ‘M’ is the subject’s mass and *W*_*co_exp*_ represents collision dissipation work. We analyze the COM power component trajectories derived from empirical data (Hosseini-Yazdi Seyed-Saleh, 2024) to calculate the step positive, negative, push-off, and collision work values for walking speeds ranging from 0.8 m.s^−1^ to 1.6 m.s^−1^ on a smooth surface (even terrain).

The trajectories are adjusted to correct for random calibration biases, and single support phase positive (rebound) and negative (preload) work and are subsequently calculated (Hosseini-Yazdi & Bertram, 2025b). Since it is shown that mechanical energy dissipation occurs during different stance phases, we employ the concept of total step dissipation or negative work. It includes the heel-strike collision and the net single support work when it is negative (Hosseini-Yazdi & Bertram, 2025b). As such, presenting total step negative work as the *W*_*neg-total*_, COM energy recovery is calculated as:

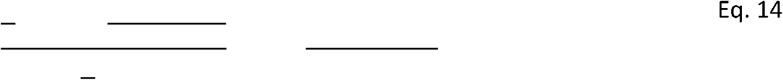

It is suggested that a portion of the step work is from storage and release of mechanical energy in tendons and other parts of the limbs (Cavagna et al., 1977; Hosseini-Yazdi Seyed-Saleh, 2024; Voloshina et al., 2013; Zelik et al., 2014). It is indicated by higher than nominal (25%) delta efficiencies. Donelan et. al. (Donelan et al., 2002a) suggested that while double support work is active, i.e. requires active muscle work, single support phase work contains some passive work, i.e. storage and release of mechanical energy. Therefore, the single support net active work is the difference between rebound and preload (Hosseini-Yazdi & Bertram, 2025b). Based on the sign of the result, either there is net positive or net negative single support active work. Accordingly, we also calculate the total step active and passive work as a fraction of total step positive work or its kinetic energy (−).

## Results

We redefined the COM energy recovery based on a step collisions approach (Donelan et al., 2002a; Ruina et al., 2005), focusing on scenarios where the COM trajectory significantly deviates from conservative inverted pendulum motion. Originally, the COM energy recovery was defined as the percentage of exchanged energy between potential and kinetic energies (Cavagna et al., 1977). Nevertheless, we assumed that the walker’s mechanical energy declined by the step dissipation. It enabled us to analytically estimate the COM energy recovery for various walking conditions using a simple powered walking model, whether the push-off was nominal or suboptimal (Kuo, 2002). Additionally, we utilized empirical data to derive our modified COM energy recovery based on heel-strike collision and total step energy loss. Subsequently, we estimate the fraction of active and passive step work.

The step length was estimated using an empirical power law for a given velocity (Kuo, 2001). As walking speed increased from 0.8 m.s^−1^ to 1.6 m.s^−1^, estimated step length increased from 0.61 m to 0.82 m (a 34.4% increase when leg length is 1 m). Correspondingly, the analytical total energy of walking (COM kinetic energy just before transition) and the push-off work during the step-to-step transition increased materially—by 400% (from 0.32 J.kg^−1^ to 1.28 J.kg^−1^) and 733.3% (from 0.03 J.kg^−1^ to 0.22 J.kg^−1^), respectively. However, the COM energy recovery declined from 90.6% to 83.1% (an 8.3% decrease). The small angle approximation yielded a recovery that was, on average, only 1.5% higher than the analytical magnitude (Figure 3).

**Figure 3.**
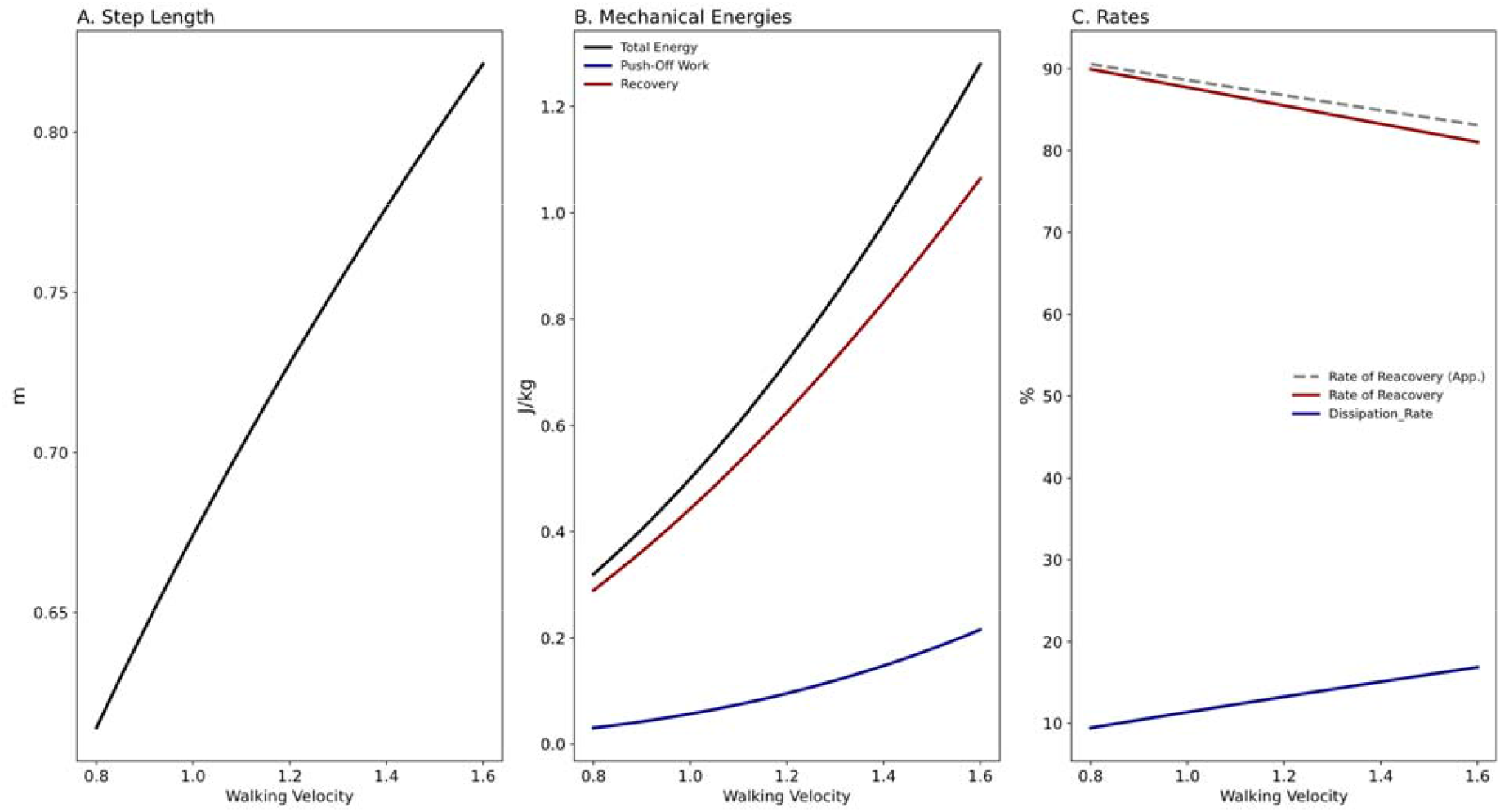
Summary of the analytic simulation for nominal walking across the given walking speed range: (A) As walking speed increases, step length also increases (leg length = 1m). (B) The total mechanical energy of the walker, push-off work, and the magnitude of mechanical energy transferred from one step to the next (recovered energy or recovery) all rise. (C) While the dissipation rate increases, the COM energy recovery declines. The dashed line represents the recovery when a small angle approximation is employed.

For suboptimal push-offs, where push-off magnitudes varied from zero to nominal (velocity = 1.25 m.s^−1^, *α* = 0.37 rad.), the mid-transition velocity angle relative to the pre-transition velocity (*γ*) increased from zero to 0.37 rad. Simultaneously, the analytical collision impulse decreased from 0.84 m.s^−1^ to 0.49 m.s^−1^ (a 71.4% reduction), while analytical push-off work increased from zero to 0.12 J.kg^−1^ and collision work declined from 0.36 J.kg^−1^ to 0.12 J.kg^−1^ (a 66.7% reduction). The analytical COM energy recovery increased from 54.5% to 84.9% (a 55.8% increase). The small angle approximation initially suggested a lower recovery but exceeded the analytical trajectory when the suboptimal push-off impulse reached approximately 70% of the optimal push-off magnitude (Figure 4).

**Figure 4.**
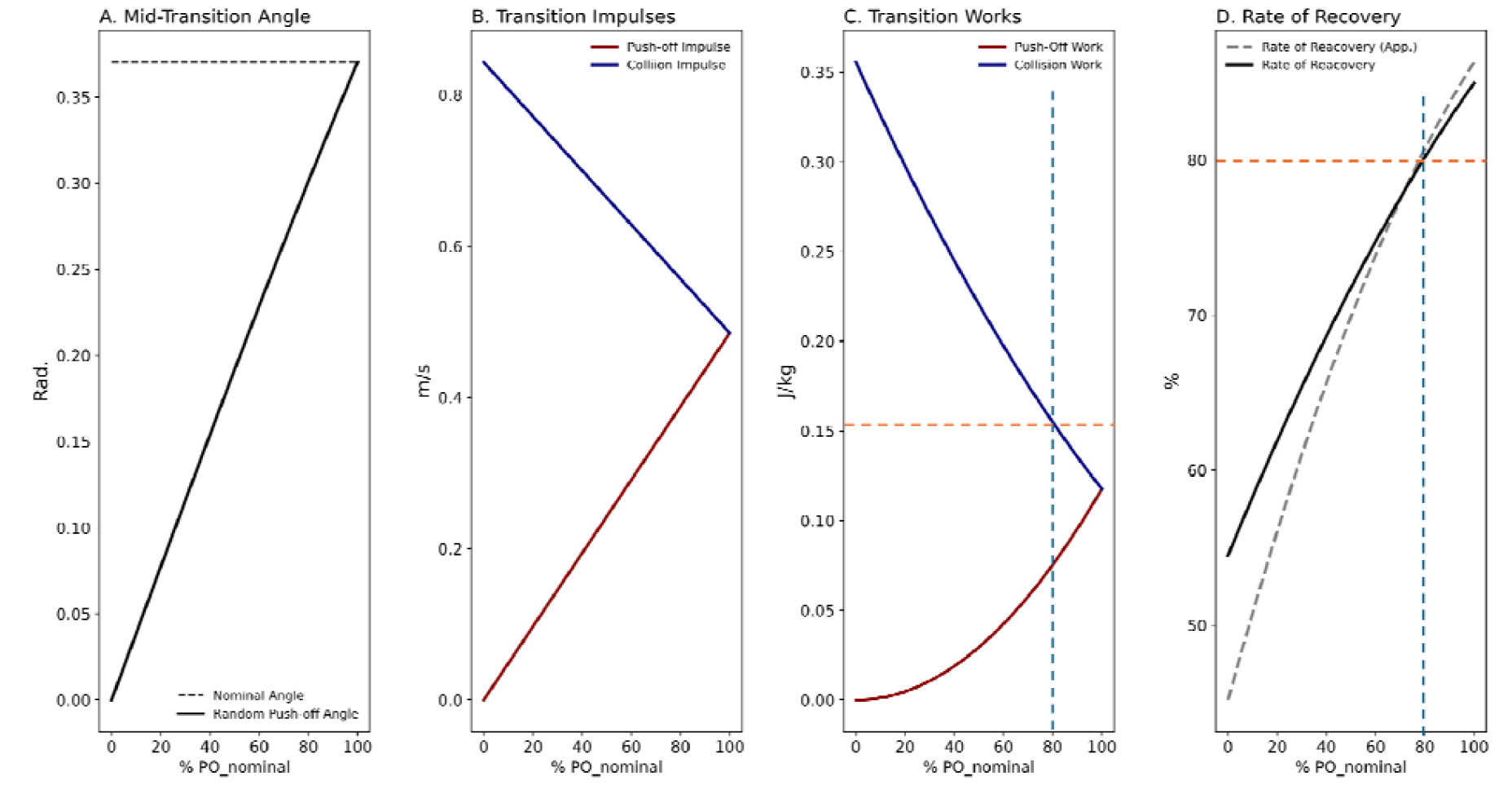
With a suboptimal push-off (), as the magnitude of the induced push-off increases (A) the mid-transition velocity angle change () rises, (B) resulting collision impulse declines, (C) the associated collision work dissipation decreases, and (D) the COM energy recovery for the suboptimal push-off elevates. The vertical dashed line depicts exerting 80% of the optimal push-off. The associated transition dissipation is 0.16 and the estimated COM energy recovery is 80%.

For a step-up, we assumed that optimal walking occurred when the entire step transition energy and the work required to overcome gravity were supplied by the pre-emptive push-off. As the step-up amplitude increased, the optimal angle of the mid-transition COM velocity change () also increased. Simulation indicated that push-off magnitudes for step-up walking exceeded those for nominal even walking at the same speed but were constrained by the maximum inducible analytical magnitude (), where the analytical collision became zero (Kuo, 2002). On average, the small angle approximation overestimated the analytical COM energy recovery by only 1.3%. The analytical COM energy recovery increased from 84.9% to 99.3% as step elevation changes varied from 0 m to 0.05 m, indicating that maximum push-off eliminated dissipation, allowing full transfer of the previous step’s mechanical energy (Figure 5).

**Figure 5.**
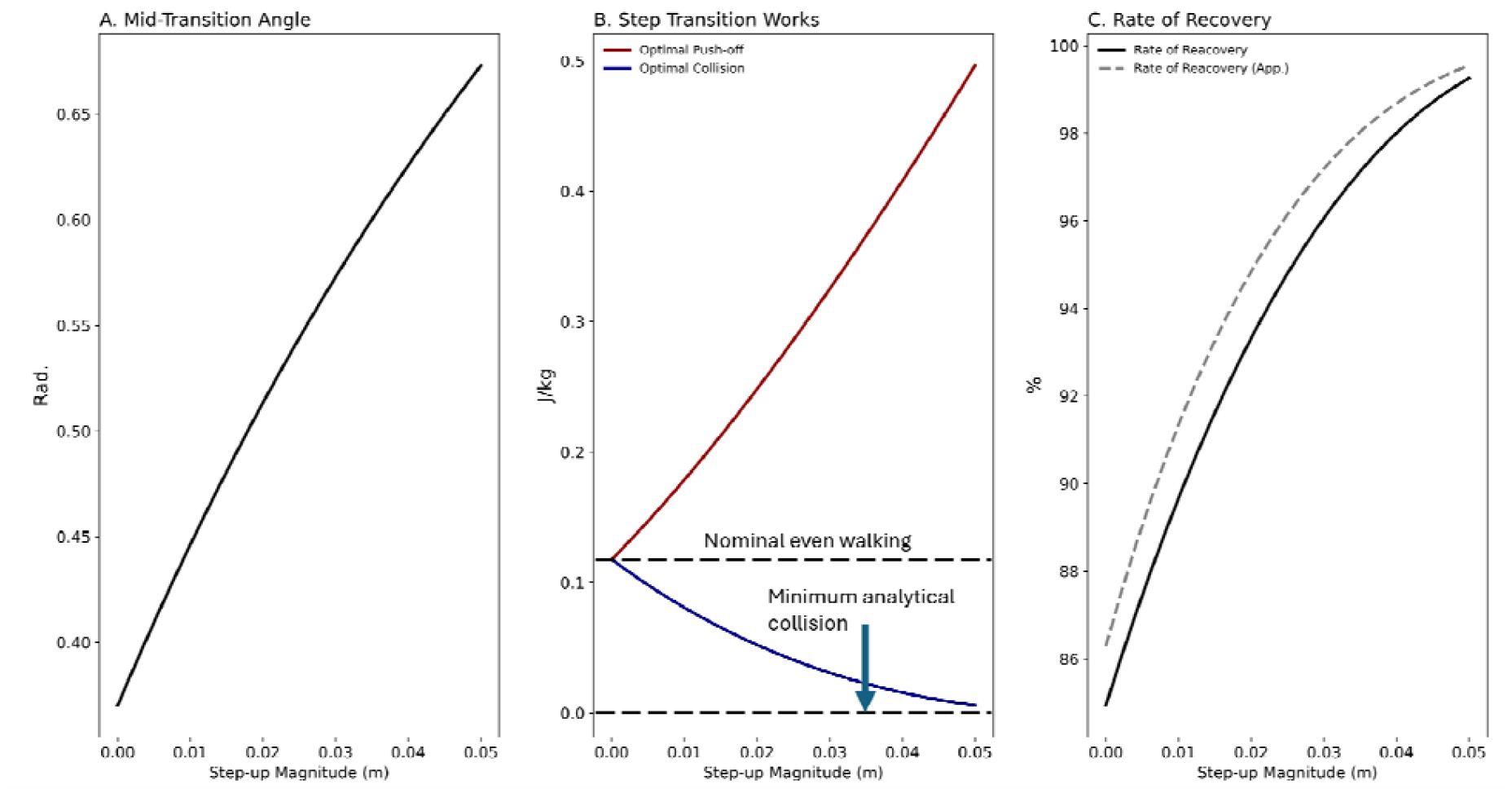
Simulation result for step-up when the entire step energy and gravity work is provided by a pre-emptive push-off: with step-up amplitude increase (A) the mid-transition velocity angle change () increases, (B) as the required push-off increases from nominal magnitude to maximum, the collision declines to zero, (C) the analytical COM energy recovery increases. The increase of COM energy recovery is because of the decrease in the heel-strike collision loss. At maximum push-off impulse, the model predicts no dissipation and as such the entire step’s energy is transferred to the next step.

For a step-down, gravitational potential energy contributes to step transition work, reducing the required push-off as its amplitude increases. Simulations showed that for = −0.036 m, no push-off was required, as gravitational potential energy fully compensated for the step transition energy, resulting in maximum collision energy dissipation. We terminated simulation when pre-emptive push-off reached zero, defining this as the maximum step-down amplitude. Beyond this threshold, the analytical collision could not offset excess gravitational energy conversion, and walkers could not sustain the average prescribed walking speed. The COM energy recovery decreased with greater step-down amplitudes, declining from 85% to 54% as collision dissipation increased from 0.12 to 0.36. For step-down, the small angle approximation overestimated the analytical recovery, but the trajectories intersected. As such, the small angle approximation underestimated the COM energy recovery at the maximum step-down amplitude (Figure 6).

**Figure 6.**
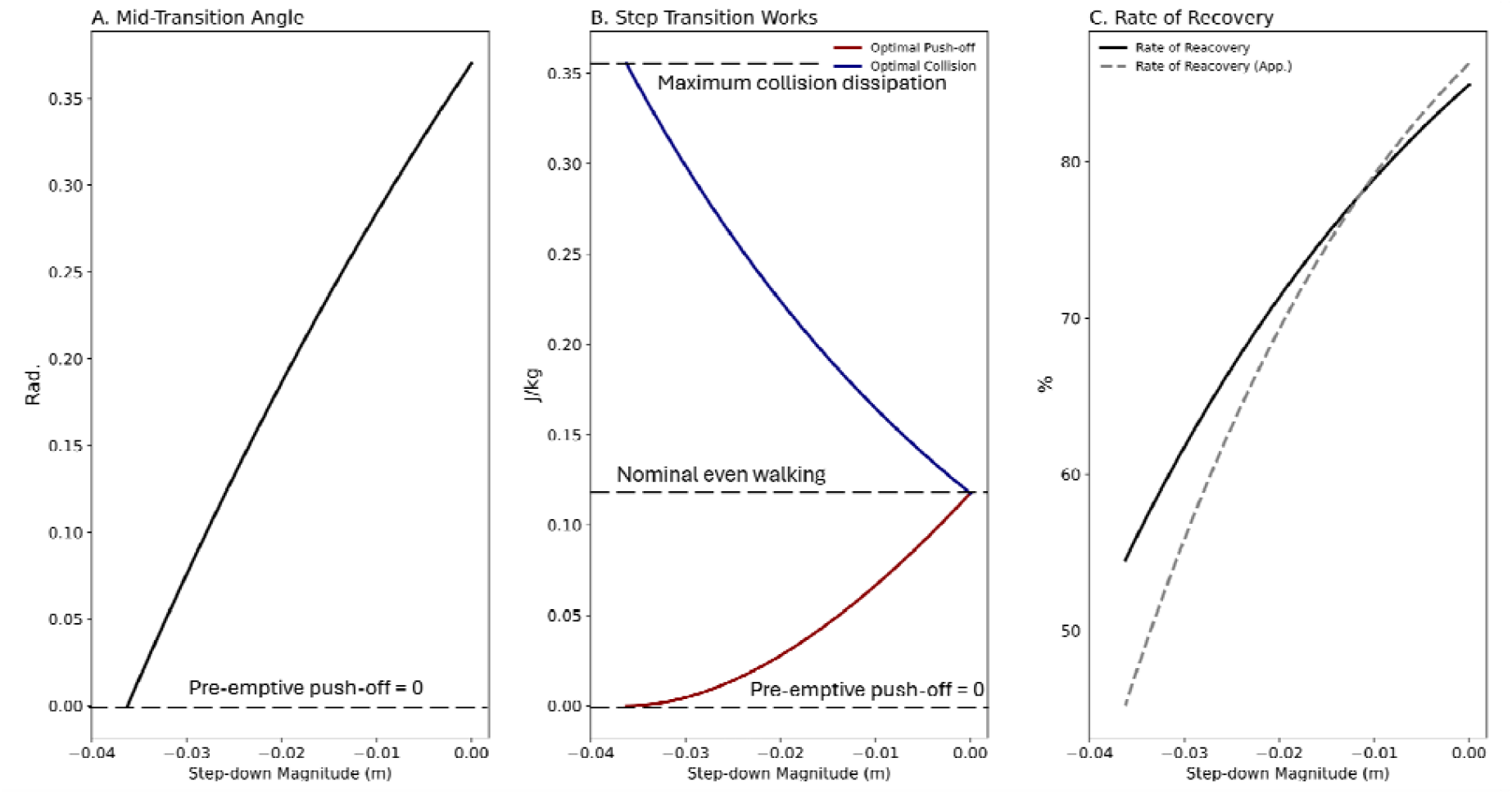
Simulation result for a step down when the entire step energy and gravity work is provided by a pre-emptive push-off and conversion of the gravity potential energy: with step-down amplitude increase (A) the mid-transition velocity angle chang () declines, (B) as the required push-off decreases from nominal magnitude to zero, the collision reaches its maximum, (C) the analytical COM energy recovery also decreases. Beyond the maximum collision impulse, the model predicts the walker’s speed increases beyond the prescribed average speed.

Using available empirical data (Hosseini-Yazdi Seyed-Saleh, 2024), based on heel-strike collision dissipation, the COM energy recovery consistently declined from 82.5% to 51.5%, closely mirroring model predictions but at a substantially faster rate. On average, with restricted lookahead, the COM energy recovery was 3.4% lower than normal lookahead. When we considered total step dissipation, analyzing the COM work components revealed a walking speed threshold below which push-off work exceeded subsequent heel-strike collision work. This surplus positive energy could increase walking speed beyond what was prescribed. Since total positive and negative step work were balanced and the average prescribed speed was maintained, the extra mechanical energy post-step transition must have been dissipated during the subsequent single support phase yielding net negative work. Examining the associated single support mechanical work indicated net negative work for walking speeds below the identified threshold, i.e. rebound – preload < 0.

Beyond the threshold speed, a net mechanical energy deficit at step transition was observed, which was compensated by the associated net positive single support work post-step transition, i.e. rebound – preload > 0. Consequently, we calculated the total step dissipation by incorporating net single support negative work and heel-strike collision loss. This adjustment showed that the COM energy recovery increased for walking speeds below the threshold, rising from 48.6% to 59.4%, with the maximum observed at v = 1.211 ⍰ m.s^−1^ (threshold). Beyond this speed, the COM energy recovery declined to 51.5%. With restricted lookahead, the maximum COM energy recovery was 58.5% at v = 1.131 ⍰ m.s^−1^ (Figure 7).

**Figure 7.**
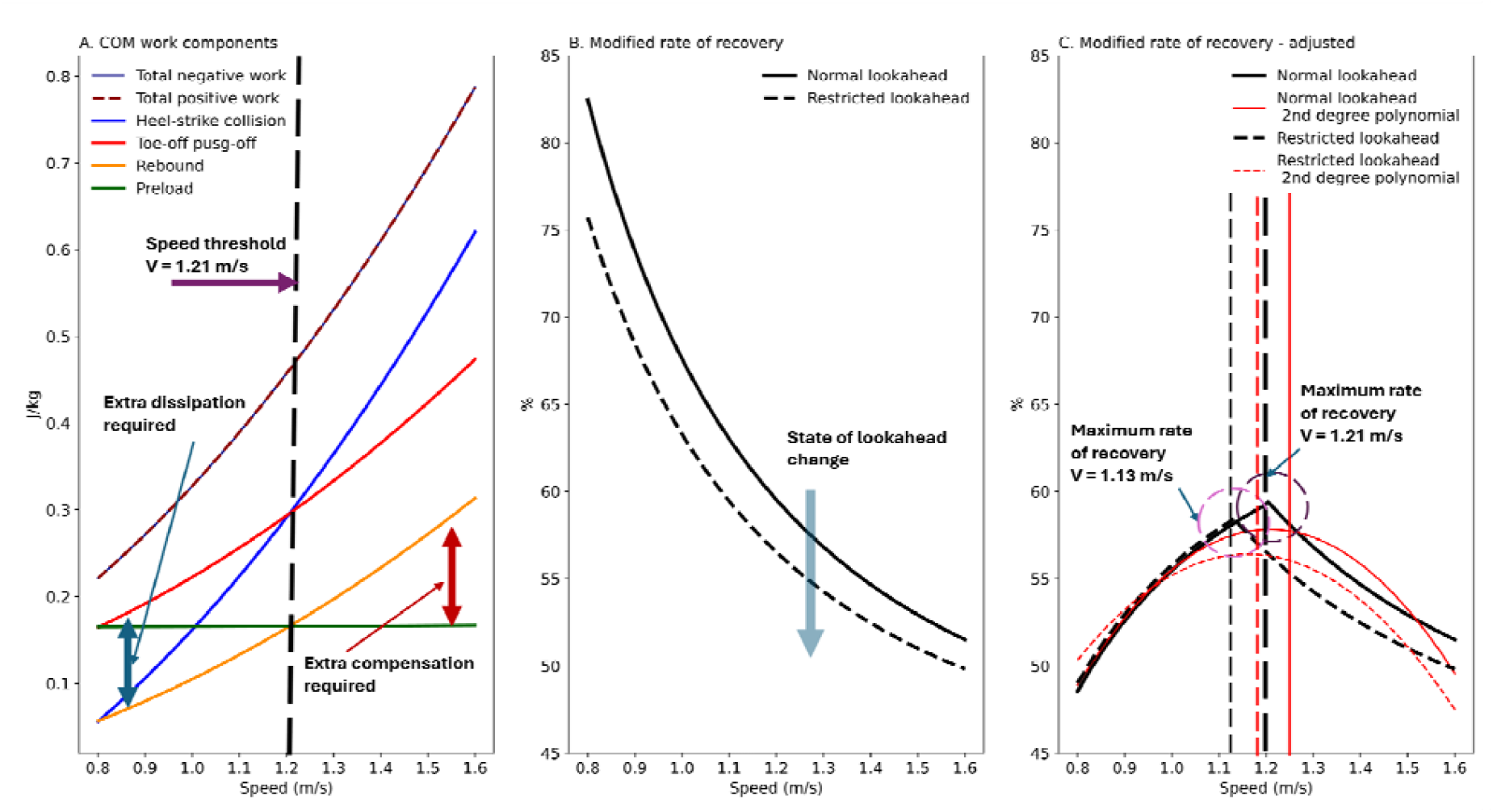
(A) The COM work components derived from empirical data (Hosseini-Yazdi Seyed-Saleh, 2024) reveal walking speeds where step transition push-off exceeds the associated step collision. This surplus mechanical energy is dissipated during the single support phase, as indicated by the net negative single support work (Rebound – Preload). (B) When only step transition collision dissipation work is considered for the rate of recovery calculation, the trend resembles that of the powered simple walking model, showing a consistent decline. With restricted lookahead, the trajectory shows more step transition dissipation than normal lookahead (C) Considering the total step dissipation as the sum of collision and net single support work when it is negative into the COM energy recovery calculation reveals an adjusted trajectory with a maximum point at v = 1.21. This speed corresponds to the point where net single support work is zero. With restricted lookahead, the maximum COM energy recovery occurs at v = 1.13. The modified definition shows that based on the total step dissipation, the trajectory of differs before and after the maximum point. We also fitted a second order polynomial to the trajectories that are indicated in red. Their associated maximum recovery are slightly higher than our modified definition, while showing rather symmetrical profiles.

For natural lookahead, at the walking speed that coincided with maximum COM energy recovery (v = 1.211⍰ m.s^−1^), the fractions of active step work were minimum; 64% of total step work and 41% of total step energy. Similarly, the fractions of passive work were maximum; 36% of total step work and 59% of total step energy. Departing from threshold speed in either direction, the fractions of active work increased while the fractions of passive work decreased (Figure 8).

**Figure 8.**
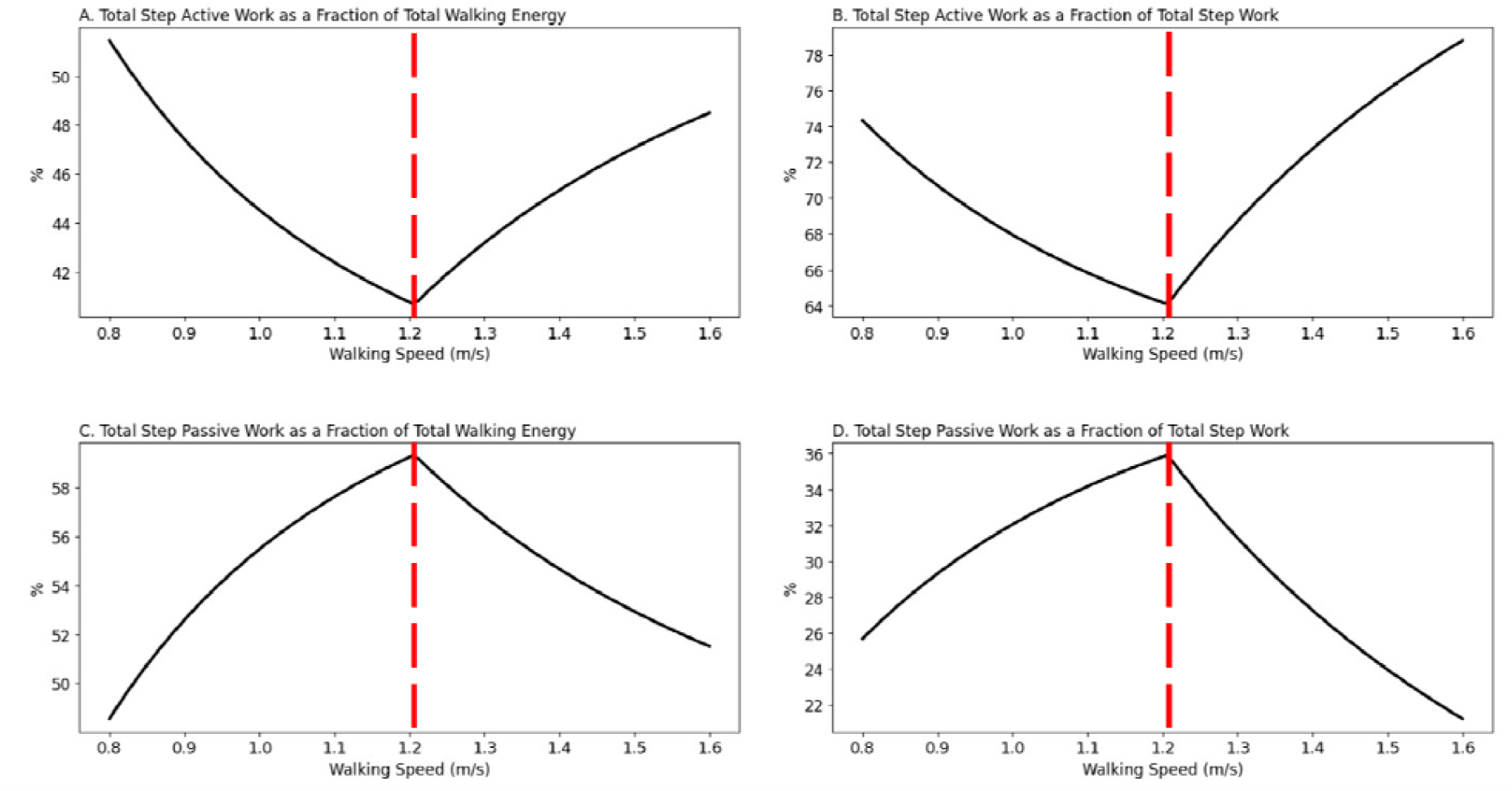
The step active (muscle work) and passive (energy release) work as fractions of total step work and total step energy for walking on an even surface with normal lookahead: (A) and (B) at the walking speed when the rate of recovery is maximum the active work fractions are minimum. (C) and (D) by the same token, the fractions of passive work are also maximum.

## Discussion

In this study, we propose a modified definition for the COM energy recovery based on total step mechanical energy dissipation (Hosseini-Yazdi & Bertram, 2025b). This metric quantifies the direct proportion of a step’s total energy that is transferred to the subsequent step. Energy dissipated during this transition is compensated by active work exerted on the COM during the stance phase (Donelan et al., 2002b; Hosseini-Yazdi & Bertram, 2025b). Hence, unlike earlier definitions which suggest that energy dissipation stems from an incomplete conversion between potential and kinetic energy (Cavagna et al., 1977), our approach attributes the step active mechanical work performance to the energy loss during the entire step phase (Bertram & Hasaneini, 2013; Hosseini-Yazdi & Bertram, 2025b; Ruina et al., 2005). This reflects the deviation from inverted pendulum motion during the step-to-step transition and COM energy modulation during single stance.

Contrary to the original definition, which posited a maximum recovery at a specific walking velocity (v ~ 1.2 or 4.3)(Cavagna et al., 1977), our analytical model reveals a nearly linear decline in COM energy recovery with increasing speed. This decrease occurs because collision impulse is directly proportional to walking velocity () (Kuo, 2002), leading to energy dissipation proportional to the square of velocity (). As a result, this method cannot predict preferred walking speeds, since the powered simple walking model (Kuo, 2002) assumes that the step transition negative and positive work are optimal and equal for any given walking speed. Therefore, the post step transition motion is completely passive. We observe similar trajectories with no optimal walking speed for other simulations. When the push-off is suboptimal, post-step transition additional active mechanical work must be performed (Hosseini-Yazdi & Bertram, 2025b; Hosseini-Yazdi & Kuo, 2025). This increases heel-strike dissipation (Hosseini-Yazdi & Bertram, 2025b), almost halving the COM energy recovery when heel-strike occurs prior to push-off. For uneven walking, the analytical COM energy recovery exhibits high variability. On completely random uneven terrain, the analytical average COM energy recovery (~82%) approximates that of even nominal walking (~85%) at a similar speed (*v*_*ave*_ = 1.25 m.s^−1^). However, uneven walking is energetically more demanding (Pandolf et al., 1976), as reflected in greater mechanical work (Hosseini-Yazdi & Kuo, 2025; Voloshina et al., 2013). During uneven walking, the proposed suboptimal timing of the pre-emptive push-off must increase the reliance on the subsequent stance-phase rebound work (Hosseini-Yazdi & Bertram, 2025c; Hosseini-Yazdi & Kuo, 2025), which leads to amplified total active mechanical work performance and thus projects a lower COM energy recovery.

The empirical COM work trajectories (Hosseini-Yazdi Seyed-Saleh, 2024) reveal an imbalance in step transition work, which is regulated during the subsequent single support phase (net single support work, Figure 7A). Consequently, we consider dissipation from both step transition (heel-strike) and the subsequent single support phase, when net work is negative, to calculate COM energy recovery. The walking speed associated with the maximum COM energy recovery closely aligns with the preferred or self-selected walking speed on even terrain (Gast et al., 2019; Hosseini-Yazdi & Bertram, 2025b; Ralston, 1958), when net single support work is zero.

As the empirical data indicates, when COM energy recovery is maximum, the net single support work is zero. A similar observation was noted by other research groups (Donelan et al., 2002a). Since the analytical model also does not assume any work for single support, the model and the total step dissipation approach must be comparable for the associated speed. However, we observe a substantial difference between model (83%) and the empirical data (59.4%) driven *R*_*c*_ when *v*_*ave*_ = 1.25 m.s^−1^. Such a difference can be associated with larger collision dissipation in human walking. In the model, the pre-emptive push-off occurs before heel-strike (Kuo, 2002). As a result, the COM velocity has completely changed (Adamczyk & Kuo, 2009) before the collision impulse. Whereas, in human walking, there is a double support phase in which both push-off and collision impulses are acting concurrently (Figure 2). Therefore, we can conclude that, before heel-strike, the push-off has not completely affected the COM velocity to limit the collision magnitude. On the other hand, there are other dissipating factors that are present during heel-strike. Some of the mechanical energy may be lost to friction, plastic deformations in shoe and foot, and some may convert to heat. Additionally, it is shown that human walking includes some negative soft tissue work (Riddick & Kuo, 2016; Van Der Zee & Kuo, 2021; Zelik & Kuo, 2010).

The lower COM energy recovery in the modified or total step dissipation method may be associated with the external work calculation approach. In the modified method, external work is calculated using a limb-by-limb approach (Donelan et al., 2002b; Kuo et al., 2005), which has been shown to yield higher work magnitudes than the combined method on which the original *R*_*IP*_ calculation is based (Cavagna et al., 1977; Donelan et al., 2002b).

One major difference of proposed COM energy recovery *R*_C_ and the original definition is when recovery = 0. In our current definition, zero recovery means mechanical energy (kinetic) loss that must be compensated by active muscle work, otherwise, the walker falls in the sagittal plane. Whereas the original definition reflects on the COM potential and kinetic energy fluctuations (Cavagna et al., 1976). Thus, a zero *R*_*IP*_ does not mean disrupted progression since any deficit is covered by muscle active work (Cavagna et al., 1977). As such, unlike the original definition, the current method is able to track mechanical energy across discrete steps, indicating the mechanisms of loss and their timing.

Although the current method and the original definition of the COM energy recovery both yield a maximum value, their trajectories differ materially. The trajectory of the original COM energy recovery may align with the best curve-fit for the data that yields a symmetrical profile around the maximum (Cavagna et al., 1976, 1977). We attain a similar trajectory when we fit second order polynomials to *R*_*C*_ derived using the total step dissipation approach on empirical data (Figure 7C). In contrast, based on total step dissipation, we suggest that the COM energy recovery varies depending on two distinct factors: the dissipation of excess mechanical energy and the compensation for mechanical energy deficits after the step-to-step transition. Consequently, the current method portrays a piecewise profile influenced by the magnitude of the pre-emptive push-off relative to the total energy required for the step transition. Such asymmetry is also presented for other gait parameters. One main example could be the Cost of Transport (COT) that is asymmetrical about the minimum point (Ralston, 1958).

Visual information about terrain is suggested to be crucial for selecting appropriate footholds (Matthis et al., 2018). It is proposed that vision about approaching terrain (lookahead) enables more efficient potential-to-kinetic energy conversions and reduces active work requirements (Hosseini-Yazdi Seyed-Saleh, 2024; Matthis et al., 2017; Matthis & Fajen, 2013). On the other hand, restricted lookahead is proposed to lead to higher collision energy loss (Hosseini-Yazdi Seyed-Saleh, 2024; Matthis & Fajen, 2013). Restricted lookahead might also increase mechanical work in the lateral direction as a function of step width (Hosseini-Yazdi Seyed-Saleh, 2024), although this component may remain small (Bauby & Kuo, 2000; Donelan et al., 2004), it further elevates required COM active work. Hence, the restricted lookahead lower maximum COM energy recovery can be associated with greater step mechanical work.

In summary, human walking may resemble inverted pendulum motion during single support that is mechanical energy conservative (Donelan et al., 2002a; Kuo et al., 2005). Nonetheless, there is significant energy loss during double support (Donelan et al., 2002a, 2002b; Kuo et al., 2005), and sometimes during the single support phase (Hosseini-Yazdi & Bertram, 2025b). These deviations result in considerable mechanical costs during or following step-to-step transitions (Donelan et al., 2002a). The current COM energy recovery emphasizes the mechanical and metabolic efficiency of a pre-emptive push-off and the contribution of single support energy modulation, highlighting their critical roles in energy conservation. Unlike the original definition of COM energy recovery (Cavagna et al., 1977, 1976), our analytical model predicts a declining trend. The analytical trajectory can be attributed to the assumption that the entire step work is performed at the step transition with inverted pendulum motion during single support. However, the empirical COM work trajectories show that this assumption holds true for only one single walking speed. In other cases, a non-zero net single support work indicates additional work performed after the step transition. Based on empirical data and the current method, we observe a walking speed at which the COM energy recovery reaches its maximum. This speed aligns closely with reported human self-selected or optimal walking speeds (Cavagna et al., 1976; Gast et al., 2019; Hosseini-Yazdi & Bertram, 2025b; Ralston, 1958). This speed also coincides with minimum active muscle work. As COM work explains the majority of walking energetics (Donelan et al., 2002a), we anticipate that any walking condition associated with elevated energetics will also show reduced COM energy recovery. Examples include walking with restricted lookahead (maximum empirical rate of recovery = 58.5% vs 59.4% for normal lookahead), on uneven terrain, or old age walking. Furthermore, the lower COM energy recovery based on empirical data may be associated with other costs or dissipation, such as soft tissue or peripheral work, that are not considered in the analytical simulation. Developing a sound metric for the COM energy recovery is particularly important for understanding gait mechanics, assessing mobility in populations such as the elderly or those with impairments, guiding rehabilitation, and optimizing prosthetic design. Additionally, *R*_*C*_ is influenced by factors like walking speed, terrain, and load, making it a valuable tool for evaluating walking performance and efficiency in both clinical and everyday settings.

## Acknowledgment

This work was supported by a Natural Sciences and Engineering Research Council of Canada (NSERC) Discovery grant (04823-2017) received by J.E.A.B.

## Notes

**Conflict of Interest:** None

### Competing Interest Statement

The authors have declared no competing interest.

### Summary of Updates

Added a new author, and also modified the process of redefining the step mechanical energy recovery to exactly address the adjustments from the original definition.

## References

Adamczyk, P. G., & Kuo, A. D. (2009). Redirection of center-of-mass velocity during the step-to-step transition of human walking. The Journal of Experimental Biology, 212(Pt 16), 2668–2678. 10.1242/jeb.027581

Alexander, R. M. (1992a). A model of bipedal locomotion on compliant legs. Philosophical Transactions of the Royal Society of London. Series B: Biological Sciences, 338(1284), 189–198.

Alexander, R. M. (1992b). Simple models of walking and jumping. Human Movement Science, 11(1–2), 3–9. 10.1016/0167-9457(92)90045-D

Alexander, R. McN. (1995). Simple Models of Human Movement. Applied Mechanics Reviews, 48(8), 461–470. 10.1115/1.3005107

Bauby, C. E., & Kuo, A. D. (2000). Active control of lateral balance in human walking. Journal of Biomechanics, 33(11), 1433–1440. 10.1016/s0021-9290(00)00101-9

Bertram, J. E. A., & Hasaneini, S. J. (2013). Neglected losses and key costs: Tracking the energetics of walking and running. Journal of Experimental Biology, 216(6), 933–938. 10.1242/jeb.078543

Cavagna, G. A., Heglund, N. C., & Taylor, C. R. (1977). Mechanical work in terrestrial locomotion: Two basic mechanisms for minimizing energy expenditure. The American Journal of Physiology, 233(5), R243–261. 10.1152/ajpregu.1977.233.5.R243

Cavagna, G. A., Thys, H., & Zamboni, A. (1976). The sources of external work in level walking and running. The Journal of Physiology, 262(3), 639–657. 10.1113/jphysiol.1976.sp011613

Darici, O., & Kuo, A. D. (2023). Humans plan for the near future to walk economically on uneven terrain. Proceedings of the National Academy of Sciences, 120(19), e2211405120. 10.1073/pnas.2211405120

Das Gupta, S., Bobbert, M. F., & Kistemaker, D. A. (2019). The Metabolic Cost of Walking in healthy young and older adults – A Systematic Review and Meta Analysis. Scientific Reports, 9(1), 9956. 10.1038/s41598-019-45602-4

Donelan, J. M., Kram, R., & Kuo, A. D. (2002a). Mechanical work for step-to-step transitions is a major determinant of the metabolic cost of human walking. The Journal of Experimental Biology, 205(Pt 23), 3717–3727. 10.1242/jeb.205.23.3717

Donelan, J. M., Kram, R., & Kuo, A. D. (2002b). Simultaneous positive and negative external mechanical work in human walking. Journal of Biomechanics, 35(1), 117–124. 10.1016/s0021-9290(01)00169-5

Donelan, J. M., Shipman, D. W., Kram, R., & Kuo, A. D. (2004). Mechanical and metabolic requirements for active lateral stabilization in human walking. Journal of Biomechanics, 37(6), 827–835. 10.1016/j.jbiomech.2003.06.002

Gast, K., Kram, R., & Riemer, R. (2019). Preferred walking speed on rough terrain; is it all about energetics? Journal of Experimental Biology, jeb.185447. 10.1242/jeb.185447

Geyer, H., Seyfarth, A., & Blickhan, R. (2006). Compliant leg behaviour explains basic dynamics of walking and running. PROCEEDINGS OF THE ROYAL SOCIETY B-BIOLOGICAL SCIENCES, 273(1603), 2861–2867. 10.1098/rspb.2006.3637

Griffin, T. M., Roberts, T. J., & Kram, R. (2003). Metabolic cost of generating muscular force in human walking: Insights from load-carrying and speed experiments. Journal of Applied Physiology, 95(1), 172–183. 10.1152/japplphysiol.00944.2002

Hosseini-Yazdi Seyed-Saleh. (2024). Energetics and Biomechanics of Uneven Walking for Young and Older Adults [University of Calgary]. https://prism.ucalgary.ca/server/api/core/bitstreams/be2fa66b-26a2-4421-9d30-dfcc55dcfb35/content

Hosseini-Yazdi, S.-S., & Bertram, J. E. (2025a). The consequence of uneven walking transitory modulation strategies: A simulation-based approach. Journal of Theoretical Biology, 614, 112234. 10.1016/j.jtbi.2025.112234

Hosseini-Yazdi, S.-S., & Bertram, J. E. A. (2025b). Center of mass work analysis predicts preferred walking speeds for varying walking conditions. Journal of Biomechanics, 185, 112682. 10.1016/j.jbiomech.2025.112682

Hosseini-Yazdi, S.-S., & Bertram, J. E. A. (2025c). Optimum Push-off for Uneven Walking Based on the Just-in-Time Strategy: Walking with Interrupted Push-Off is Mechanically Costly. Journal of Biomechanical Engineering, 1–20. 10.1115/1.4069666

Hosseini-Yazdi, S.-S., & Kuo, A. D. (2025). The energetic cost of human walking as a function of uneven terrain amplitude. Journal of Experimental Biology, jeb.249840. 10.1242/jeb.249840

Huang, T. P., Shorter, K. A., Adamczyk, P. G., & Kuo, A. D. (2015). Mechanical and energetic consequences of reduced ankle plantarflexion in human walking. Journal of Experimental Biology. 10.1242/jeb.113910

Kuo, A. D. (2001). A simple model of bipedal walking predicts the preferred speed-step length relationship. Journal of Biomechanical Engineering, 123(3), 264–269. 10.1115/1.1372322

Kuo, A. D. (2002). Energetics of Actively Powered Locomotion Using the Simplest Walking Model. Journal of Biomechanical Engineering, 124(1), 113–120. 10.1115/1.1427703

Kuo, A. D., Donelan, J. M., & Ruina, A. (2005). Energetic Consequences of Walking Like an Inverted Pendulum: Step-to-Step Transitions: Exercise and Sport Sciences Reviews, 33(2), 88–97. 10.1097/00003677-200504000-00006

Matthis, J. S., Barton, S. L., & Fajen, B. R. (2017). The critical phase for visual control of human walking over complex terrain. Proceedings of the National Academy of Sciences, 114(32). 10.1073/pnas.1611699114

Matthis, J. S., & Fajen, B. R. (2013). Humans exploit the biomechanics of bipedal gait during visually guided walking over complex terrain. Proceedings of the Royal Society B: Biological Sciences, 280(1762), 20130700. 10.1098/rspb.2013.0700

Matthis, J. S., Yates, J. L., & Hayhoe, M. M. (2018). Gaze and the Control of Foot Placement When Walking in Natural Terrain. Current Biology, 28(8), 1224–1233.e5. 10.1016/j.cub.2018.03.008

Pandolf, K. B., Haisman, M. F., & Goldman, R. F. (1976). Metabolic energy expenditure and terrain coefficients for walking on snow. Ergonomics, 19(6), 683–690. 10.1080/00140137608931583

Ralston, H. J. (1958). Energy-speed relation and optimal speed during level walking. Internationale Zeitschrift Für Angewandte Physiologie Einschliesslich Arbeitsphysiologie, 17(4), 277–283. 10.1007/BF00698754

Riddick, R. C., & Kuo, A. D. (2016). Soft tissues store and return mechanical energy in human running. Journal of Biomechanics, 49(3), 436–441. 10.1016/j.jbiomech.2016.01.001

Ruina, A., Bertram, J. E. A., & Srinivasan, M. (2005). A collisional model of the energetic cost of support work qualitatively explains leg sequencing in walking and galloping, pseudo-elastic leg behavior in running and the walk-to-run transition. Journal of Theoretical Biology, 237(2), 170–192. 10.1016/j.jtbi.2005.04.004

Van Der Zee, T. J., & Kuo, A. D. (2021). Soft tissue deformations explain most of the mechanical work variations of human walking. Journal of Experimental Biology, 224(18), jeb239889. 10.1242/jeb.239889

Voloshina, A. S., Kuo, A. D., Daley, M. A., & Ferris, D. P. (2013). Biomechanics and energetics of walking on uneven terrain. Journal of Experimental Biology, jeb.081711. 10.1242/jeb.081711

Zelik, K. E., Huang, T.-W. P., Adamczyk, P. G., & Kuo, A. D. (2014). The role of series ankle elasticity in bipedal walking. Journal of Theoretical Biology, 346, 75–85. 10.1016/j.jtbi.2013.12.014

Zelik, K. E., & Kuo, A. D. (2010). Human walking isn’t all hard work: Evidence of soft tissue contributions to energy dissipation and return. Journal of Experimental Biology, 213(24), 4257–4264. 10.1242/jeb.044297

